# Measuring technical variability in Illumina DNA methylation microarrays

**DOI:** 10.1101/2023.11.28.569087

**Authors:** Anderson A. Butler, Jason Kras, Karolina Chwalek, Enrique I. Ramos, Isaac Bishof, David Vogel, Daniel L. Vera

**Affiliations:** VoLo Foundation, Jupiter, FL 33468; Voloridge Health, Jupiter, FL 33477

## Abstract

DNA methylation microarrays have become a widely used tool for investigating epigenetic modifications in various aspects of biomedical research. However, technical variability in methylation data poses challenges for downstream applications such as predictive modeling of health and disease. In this study, we measure the impact of common sources of technical variability in Illumina DNA methylation microarray data, with a specific focus on positional biases inherent within the microarray technology. By utilizing a dataset comprised of multiple, highly similar technical replicates, we identified a chamber number bias, with different chambers of the microarray exhibiting systematic differences in fluorescence intensities and their derived methylation beta values, which are only partially corrected for by existing preprocessing methods, and demonstrate that this positional bias can lead to false positive results during differential methylation testing. Additionally, our investigation identified outliers in low-level fluorescence data which might play a role in contributing to predictive error in computational models of health-relevant traits such as age.

## Introduction

In recent years, DNA methylation (DNAm) has received considerable interest as a potential biomarker for numerous clinically relevant traits, including disease and health risks^1,2^. In oncology, DNAm microarray-based assays which predict tumor site of origin have begun to see approved clinical use in the US and European Union for diagnostic purposes and for classifying cancers of unknown primary origin^3,4^, and numerous methylation-derived risk scores (MRS) and predictive models for traits such as smoking status, body weight, and age are well-established in the scientific literature^3^.

Improvements to analytical pipelines for array-based measurement of DNAm remain a high priority within the field, as technical limitations to data quality and consistency across batches currently interfere with the reliable detection of differences in DNAm, limiting the feasibility of DNAm microarray technologies for both research and clinical applications^3,5^. There is a general agreement regarding the need to account for batch and positional effects and other technical artifacts which influence microarray data and might otherwise result in spurious relationships^3^. To this end, many preprocessing methods have been developed which reduce the impact of positional effects and improve reliability. Several systematic studies have been undertaken to describe the efficacy and suitability of these methods in varying use cases. However, these studies are reliant on *in silico* experiments conducted either on repurposed experimental data or on simulated datasets. While highly valuable, conclusions drawn from such datasets must be interpreted with caution, given the potential for confounding latent variables in experimental data and potential discrepancies between simulated and real-world data.

Previous studies have examined the scale of technical noise within microarray experiments, primarily by measuring the intra-class correlation (ICC) as a metric of reliability for microarray probes. As the use of technical duplicate samples is optimal for studies aiming to measure ICC^6,7^, these studies have limited applicability to those seeking to understand technical variability within microarrays^8^.

While useful and informative, the wider utilization of established ICCs is limited by the metric itself, which measures technical variability only as it relates to biological variability for each CpG. However, as several authors examining the metric have noted, CpGs with presumably small levels of technical variability are often reported as unreliable due to the very low levels of biological variability for many CpGs within a single study and tissue type^8–11^. As ICC for any given CpG might be expected to increase, perhaps dramatically, in studies examining multiple tissue types, populations, or importantly, disease states^12^, we proposed here to explore the technical variability of Illumina microarray technology with an experimental approach that allows for the measurement of sources of technical variability without regard for biological variability.

Here, we characterized technical variability in Illumina’s MethylationEPIC v1 microarray with an experimental design which allows us to isolate and measure the effect of multiple sources of technical variability. We directly examine the contribution of common positional effects to technical variability in methylation beta values within a set of highly similar replicates using linear mixed effects models and assess the performance of preprocessing in correcting for measurement differences across technical replicates. We explore the implications of our findings regarding technical variability and alternative preprocessing methods in the context of their application to experimental design, by examining the introduction and elimination of false positives as well as the impact of preprocessing on detectable biological variability with the data. Further, we examine the impact of preprocessing on well-accepted modeling targets in the literature: epigenetic age clocks.

## Results

### Experimental design to measure technical variability in Illumina’s MethylationEPIC microarrays

We first sought to design an experiment that would allow a comparison of positional effects in Illumina’s DNA methylation microarrays. We concentrated on isolating the chamber numbers (commonly known as Sentrix Positions) and slides (Sentrix Barcodes) to measure the contribution of positional effects to the variability observed among technical replicates.

For each of four human subjects, a single vial of venous whole blood was collected and dispersed into aliquots prior to short-term frozen storage. Subsequently, eight of the aliquots for each subject were processed to isolate DNA for hybridization to microarrays, and then further split into two aliquots prior to deamination, amplification, and fragmentation (Fig 1A). A total of eight samples, one from each of the DNA extractions, were then combined to form a pooled sample for each subject. Finally, eight aliquots from each subject’s pool and eight independently isolated technical replicates for each subject were profiled using Illumina EPIC v1 microarrays, for a total of 16 assays per subject.

**Figure 1.**
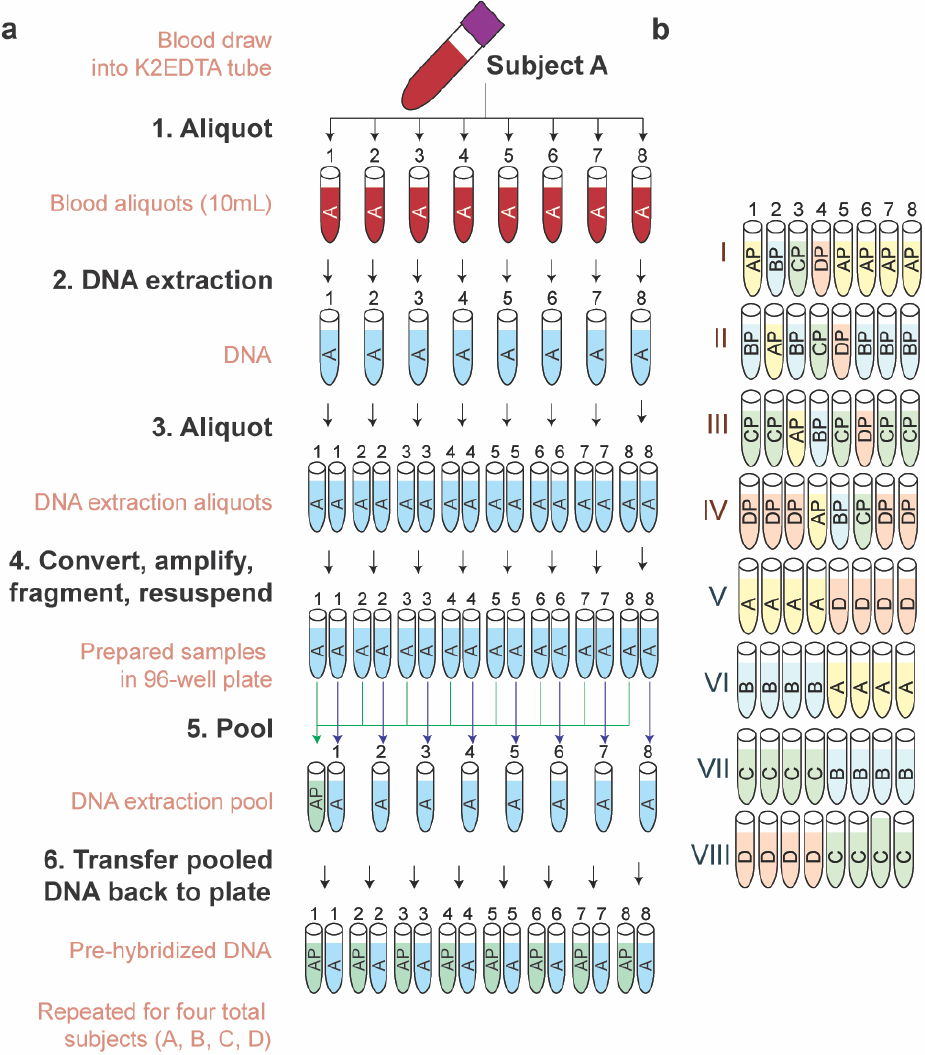
Schematic of experimental design to measure stability of microarray experiments. **(A)** Schematic of experimental design. Briefly, for each human subject, DNA was isolated from eight replicate blood aliquots, and then independently bisulfite converted, amplified, and fragmented. An aliquot of DNA from each of the eight replicates was combined into a pooled DNA sample. Both independently isolated and pooled technical replicates were measured across using Infinium MethylationEPIC microarrays. **(B)** Layout of pooled and independently prepared samples from each subject (Letters) with regard to chamber number/Sentrix position (numbers) within each slide/Sentrix Barcode (Roman numerals). Samples were used for independent statistical assessments as indicated.

To ensure that all positional effects were assayed, we measured each pooled replicate once in every chamber number, and at least once on each of four slides. To further examine the relative impact of sample preparation on variability, we also assayed eight independently prepared (unpooled) technical replicates for each subject, bring the total number of technical replicates per subject to 16 (Fig 1B). These additional samples were organized for the purposes of examining the impact of positional effects on differential methylation testing.

### Multivariate Analysis and Statistical Comparison of Positional Effects on CpGs in the Illumina MethylationEPIC platform

To begin to identify the main sources of variation within the Illumina MethylationEPIC dataset, we first conducted a principal component analysis (PCA) on beta values generated with the widely used preprocessing tool ‘SeSAMe’, using the authors’ recommended settings. Using the first fifty principal components as input, we performed hierarchical clustering of technical replicates. As each of the 16 total samples for each human subject were derived from a single blood draw (8 pooled plus 8 independently processed technical replicates), with the degrees of difference between them solely dependent upon technical factors, we anticipated that replicates from the same subject would cluster together, with biological variation between them as the main source of variation. Surprisingly, while several clusters appeared to be segregated by subject, one cluster contained intermingled samples from multiple subjects, suggesting that technical noise in methylation data may obfuscate biological signal (Fig 2A). This lack of separation by subject is notable within the first two principal components (Fig 2B). Importantly, we noted stratification by chamber number, but not slide number, within the first two principal components (Fig 2C-D).

**Figure 2.**
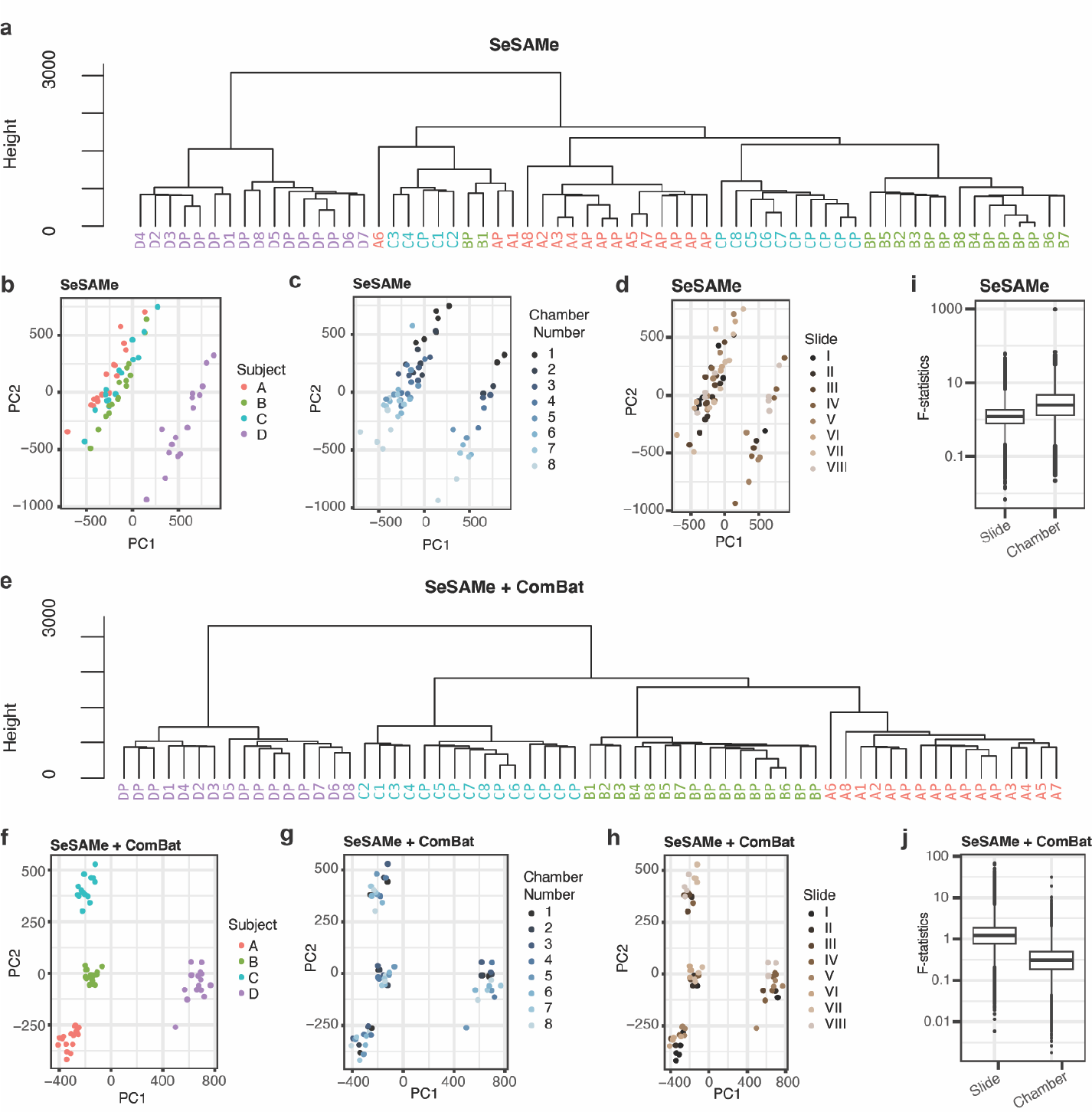
Multivariate Analysis and Statistical Comparison of Positional Effects on CpGs in the Illumina MethylationEPIC platform. **(A)** Hierarchical clustering dendrogram representing the clustering of subjects based on the first 50 principal components from methylation beta values after preprocessing with SeSAMe. The vertical axis represents the dissimilarity between subjects, while the horizontal axis shows the subjects’ labels. **(B)** Principal component analysis (PCA) plot, representing the first principal component (PC1) against the second principal component (PC2) for each sample. Each data point represents an individual technical replicate, with colors indicating different subjects. **(C-D)** PCA plots showing the first two principal components for each sample, with colors indicating different chamber numbers **(C)** or different slides **(D). (E)** Hierarchical clustering dendrogram representing the clustering of subjects based on the first 50 principal components from adjusted beta values after preprocessing with SeSAMe and ComBat adjustments for chamber number. **(F-H)** PCA plots showing clustering of subjects after ComBat adjustment for chamber number, colored by subject **(F)**, chamber number **(G)**, or array **(H). (I-J)** Box plots presenting F statistics obtained from analysis of variance (ANOVA) performed on beta values preprocessed either with SeSAMe alone **(I)** or with SeSAMe + ComBat using chamber number as batch **(J)**. The box represents the interquartile range (IQR), with the median indicated by a line inside the box. Whiskers extend to the minimum and maximum values within 1.5 times the IQR. Data points beyond 1.5 times the IQR are plotted individually.

Given the observed stratification, we hypothesized that some bias in the data had been introduced by positional effects, particularly chamber number. To explore this hypothesis, we used ComBat, a commonly used empirical Bayes method for removing technical variation in DNA methylation microarray data, to adjust DNA methylation beta values for either chamber number or slide number. After controlling for slide, we noted no improvement in segregation in hierarchical clustering (Fig S1A) or within the first two principal components (Fig S1B-D); rather, clustering appeared worsened, with subject D less cleanly segregated than in unadjusted data. In contrast, after controlling for chamber number, we noted a perfect segregation by subject identity (Fig 2G), as well as improved biological signal within the first two principal components (Fig 2H).

To gain a greater understanding of how each of the positional effects impacts variability within the Illumina MethylationEPIC microarray, we examined the contribution of these effects to variance in methylation beta values using linear mixed effects models.

For each CpG, we used linear mixed effects regression to estimate the proportion of variance in beta values explained by the chamber and slide number positional effects. To compare aggregate data from all CpGs, we recorded the F-statistic as a metric of the impact of these positional effects. We observed that in data preprocessed with SeSaMe, the explainable variance due to chamber number is larger than the effect of slide (Fig 2I). Consistent with this observation, correcting for the chamber number with ComBat appears to result in a greater overall reduction in explainable noise than correcting for the effect of slide, at least in the present dataset (Fig 2J & Fig S1E).

### Differential methylation testing on alternatively preprocessed methylation beta values from technical replicates

Having examined the contributions of positional effects to principal components of beta values, we next sought to determine the degree to which preprocessing pipelines might reduce beta value variability from a practical standpoint.

To accomplish this, for each CpG, we examined the ratio of the standard deviation (SD) between same-subject technical replicates to the SD for all replicates within the study. While replicates originating from the same initial sample are expected to have minimal biological variability, blood samples are also expected to have a high degree of similarity across subjects. Thus, to account for CpGs with very low biological variability, we added a small offset (0.0001) to the denominator when calculating SD ratios.

We observed a marked reduction in SD ratio after using the authors’ recommended settings for SeSAMe, and a further reduction after correcting for the chamber number positional effect with either ComBat on beta values, or by correcting SeSAMe-adjusted fluorescence intensities using the ComBat-Seq algorithm (Fig 3A).

**Figure 3.**
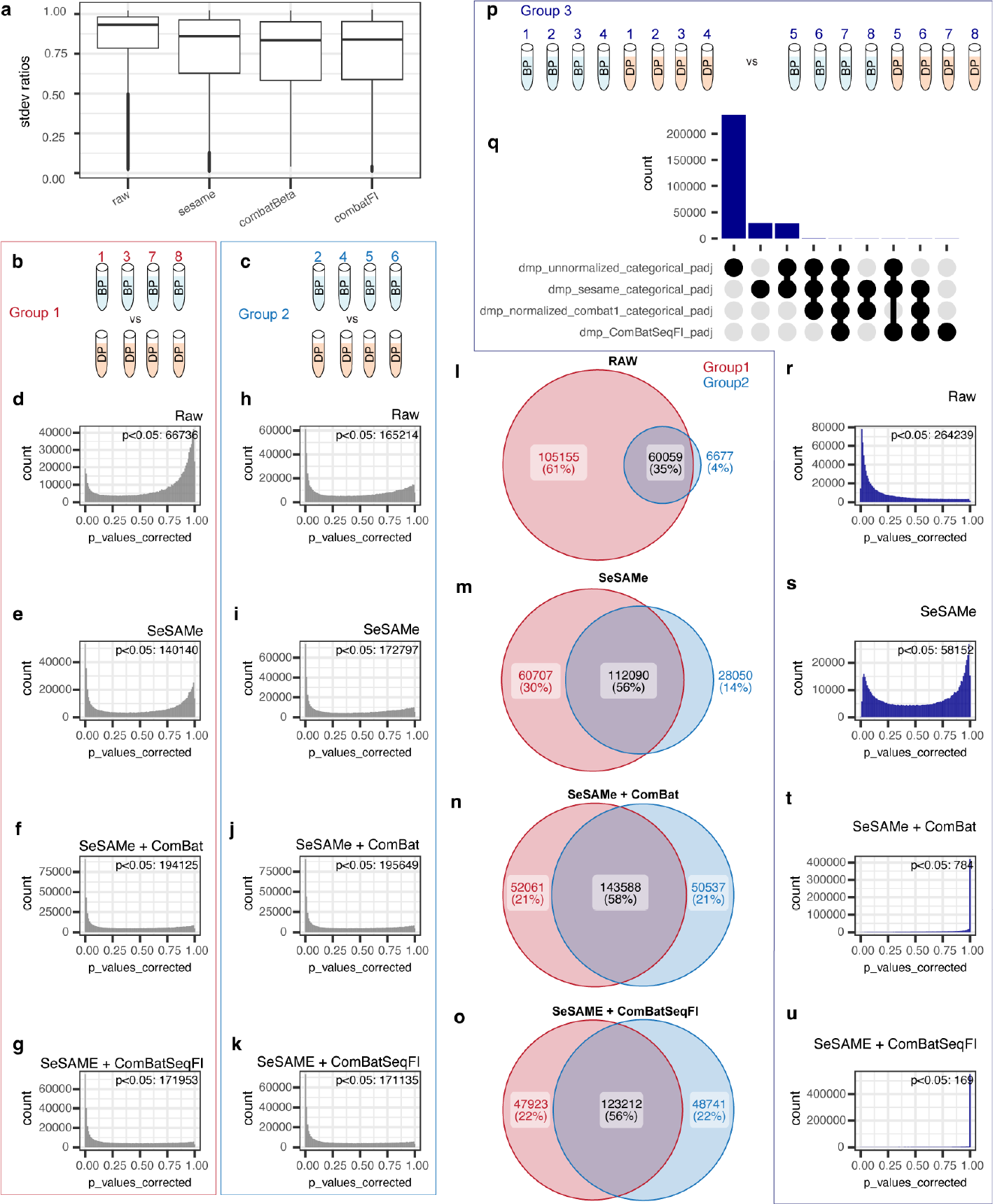
Differential methylation analysis of alternatively preprocessed methylation beta values from technical replicates. **(A)** Ratio of averaged standard deviation within subjects to standard deviation across all subjects for 614,831 CpGs shared across all preprocessing pipelines shown. **(B-C)** Representation of samples used for differential methylation testing in comparison group 1. Experimental subjects B and D were each represented by n=4 technical replicates, with each replicate located on a different array for comparison group 1 **(B**) and each replicate on the same array for group 2 **(C). (D-K)** Histograms depicting BH adjusted p-values for all CpGs after differential methylation testing of beta values using the specified preprocessing pipeline for group 1 (in red box) and group 2 samples (in light blue box). Number of CpGs with BH-adjusted p-values < 0.05 listed on upper right of each plot. **(L-O)** Venn diagrams representing the number of CpGs detected as significantly different between subjects B and D in Group 1 (red circle), Group 2 (blue circle), or in both sets of samples (overlap), using the specified preprocessing pipeline. **(P)** Representation of samples used for differential methylation testing to screen for the introduction of false positives due to positional effects. **(Q)** Upset plot relating the number of false positives within and between preprocessing pipelines (limited to 9-most abundant comparisons). **(R-U)** Histograms depicting BH adjusted p-values for all CpGs after differential methylation testing of beta values using the specified preprocessing pipeline. Number of CpGs with BH-adjusted p-values < 0.05 listed on upper right of each plot.

There are reports in the literature that preprocessing pipelines, and in particular correcting for positional effects using the ComBat algorithm, might obfuscate biological variability or introduce false positives^13^. Therefore, we next sought to examine whether this was the case in our dataset. To accomplish this, we used the dmpFinder function from the minfi R package to conduct a series of differential methylation tests on subsets of samples (Fig 3B-K).

As our data do not contain any positive controls, rather than test for false negatives, we instead examined the consistency between two independent differential methylation tests on groups of biologically different technical replicates (testing groups 1 & 2) as shown (Fig 3B-C). As testing groups 1 & 2 are comprised of pooled technical replicates from the same two subjects, we are assured that any differences between the results of the two tests arise from technical noise rather than biological differences.

If preprocessing pipelines were masking biological signal, we would expect that the total number of differentially methylated CpGs observed would decrease relative to the number observed in the raw data. However, instead we observed that in both testing groups 1 & 2 the number of differentially methylated CpGs increases after preprocessing with SeSAMe, and is further increased by adjusting the beta values for positional effects with ComBat (Fig 3D-K). Similarly, we find that the agreement between results is markedly improved with preprocessing using SeSAMe, and further improved by adjusting the beta values for positional effects with ComBat (Fig 3L-O).

Next, we sought to examine whether preprocessing might introduce false positives as previously reported^13^. To test this hypothesis, we examined a third differential methylation test, in which technical replicates from two biological samples were equally represented across the groups being compared. As the underlying biological makeup across the comparison is identical, we assume that any statistically significant results from the analysis must be falsely positive.

We observed that large many sites appear to be erroneously identified as differentially methylated when examining tests on raw or SeSAMe-adjusted beta values (Fig 3Q-S). While the number of false positives is improved after preprocessing using SeSaMe, many false positive results remain until correction for chamber number via ComBat (Fig 3T). Interestingly, the greatest reduction in false positives in this assessment was achieved using the ComBat-seq algorithm^14^, which we adapted to correct for positional effect on SeSAMe-adjusted fluorescence intensities (Fig 3U).

Together, these results suggest that from a practical standpoint, the preprocessing pipelines tested here appear to reduce beta value variability and reduce the number of false positive results, while also preserving, or potentially unmasking, biological variability.

### Fluorescence and beta values are biased by positional effects in technical replicates

Having established that chamber number corresponds to a large technical bias in beta values, we next sought to understand the origins of this bias. We first assessed low-level fluorescence data from each sample for differences in intensity. As the mean and variance for both fluorescence intensities and beta values might be expected to differ between subjects for the same CpGs, we first centered and scaled values within technical replicates from the same subject, then averaged these centered and scaled values across four observations from different subjects as shown (Fig 4A). To allow for a direct comparison between color channels, we limited our analysis of raw fluorescent intensities to Type II probes, in which the methylated and unmethylated signals are recorded on the same bead in the green and red channels, respectively. We noted a marked difference in relative signal between chamber numbers in both the green and red fluorescent channels that in raw data, with fluorescence values recorded in chambers 1 and 2 appearing less bright than other chambers. Additionally, there were apparent differences in the distributions of values between the color channels, with chamber 1 appear right-skewed while some others appeared to have varying degrees of heteroskedasticity (Fig 4B-C).

**Figure 4.**
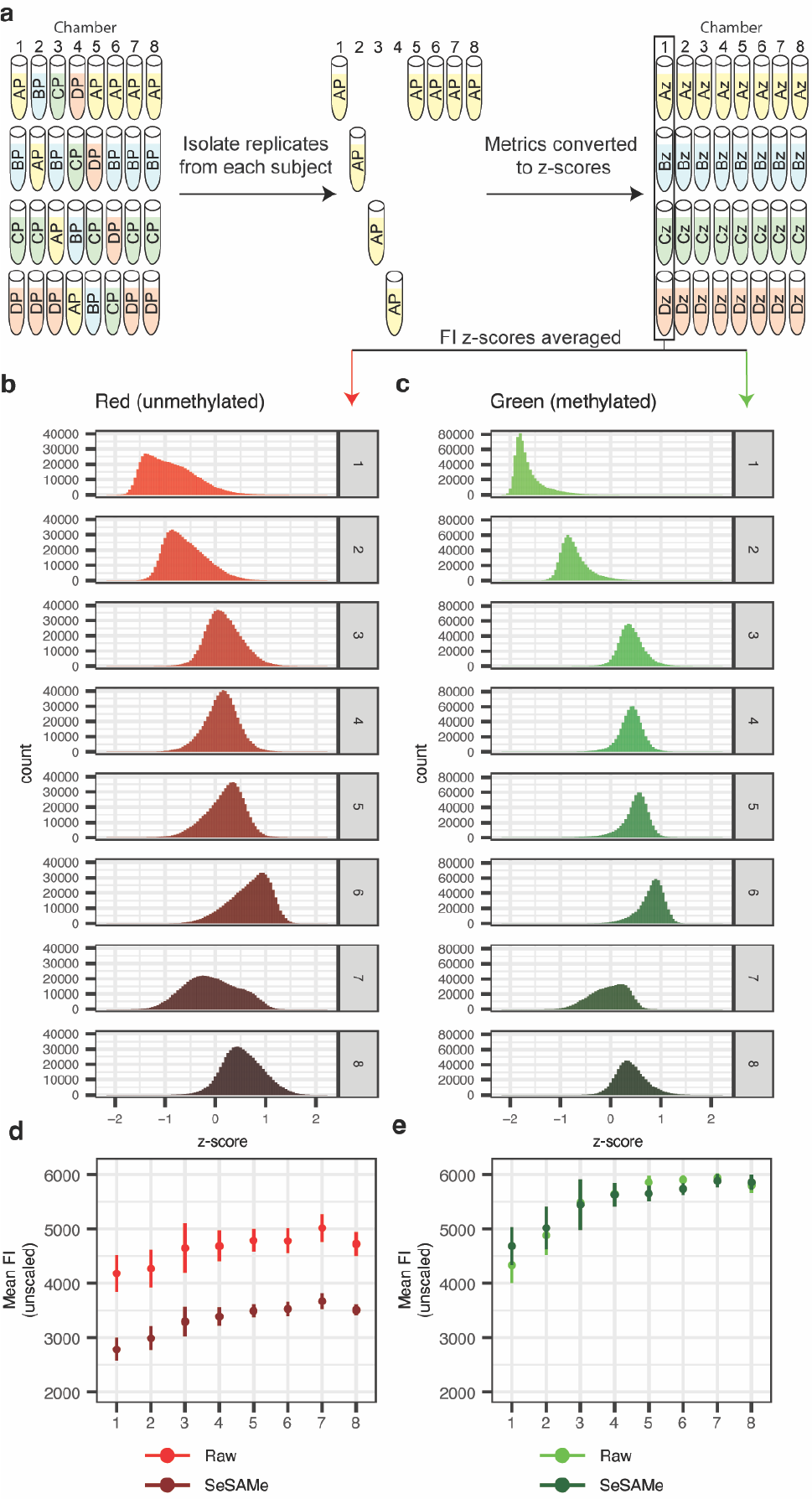
Fluorescence values compared in technical replicates measured in different chamber numbers. **(A)** Configuration of samples within slides/chambers for analysis in panels c-d. The average of four within-subject centered and scaled fluorescence intensity values for each CpG/chamber number were extracted from slides, as depicted. **(B-C)** Distribution histograms illustrating within-subject centered and scaled fluorescence intensities (FI) for all Type II probes on the MethylationEPIC array in the red **(B)** and green **(C)** channels. **(D-E)** Mean and standard deviation of fluorescence intensities (not centered or scaled) from all type 2 probes within in each chamber number.

While differences between chambers might appear large when scaled, it might be possible that chamber number biases are magnified by the scaling process, as many probes have very low variability. To ascertain whether this might be the case, we averaged fluorescence intensities of all probes in the red/green channels for each sample, and then averaged these sample means across all samples sharing the same chamber number, without centering or scaling. Similar to the results described above, we observed that the averaged mean fluorescence intensity in chamber number 1 was lower than in the other chambers, in both the red and green channels. Importantly, this bias in fluorescence intensities remained in both the green and red channel after preprocessing the data with SeSAMe (Fig 4D-4E).

We next sought to examine whether this chamber number bias impacted beta values, and indeed upon examining the averaged centered and scaled beta values from technical replicates, we observed a similar pattern as above, with technical replicates in chamber 1 and 2 appearing to show lower methylation overall than those in other chambers (Fig 5A).

**Figure 5:**
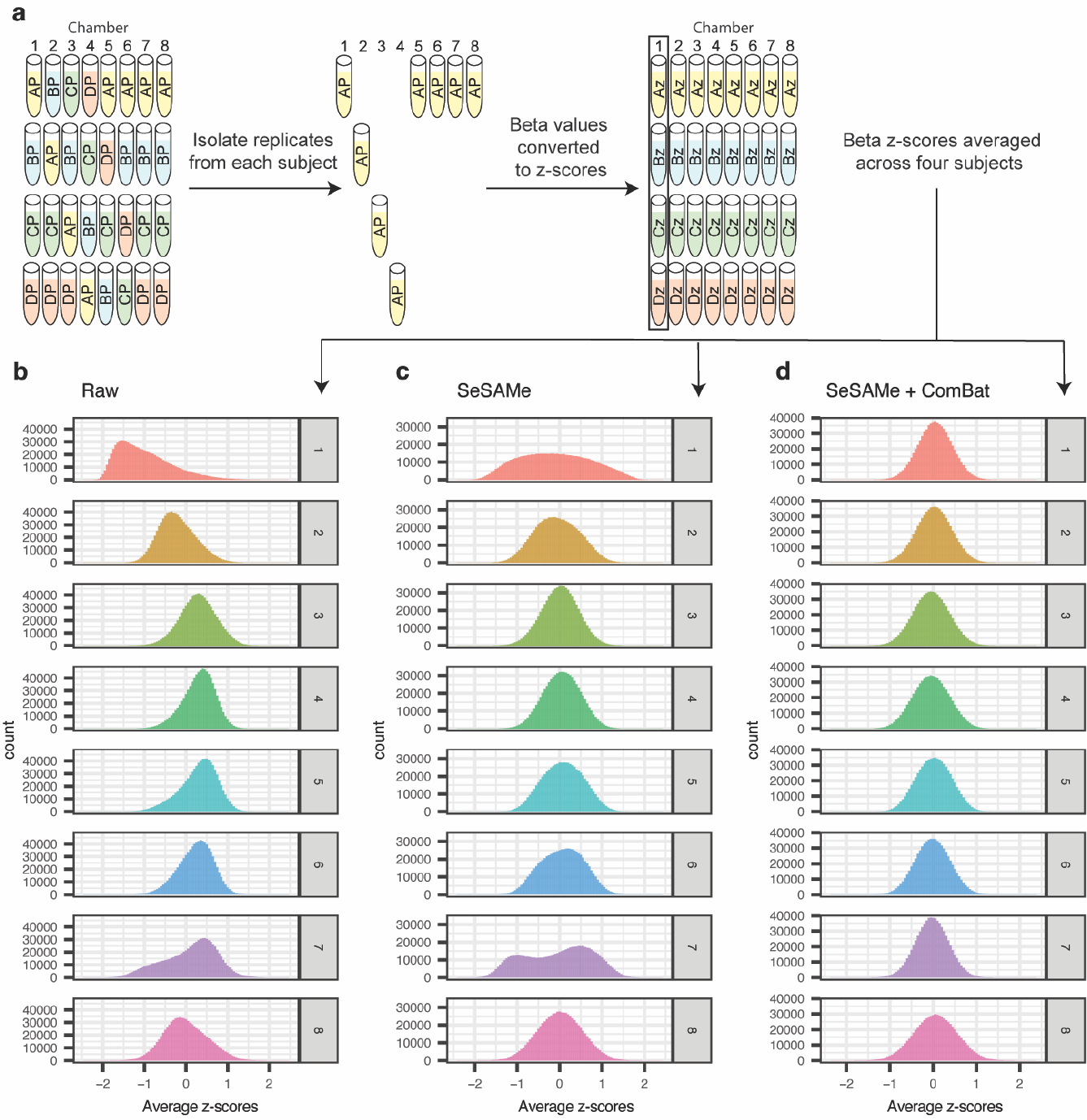
Beta values compared in technical replicates measured in different chamber numbers. **(A)** Configuration of samples within slides/chambers. The average of four within-subject centered and scaled beta values for each CpG/chamber number were extracted from slides, as depicted. **(B-D)** Histograms representing the distributions of within-subject centered and scaled beta values for all probes on the MethylationEPIC array. The histograms correspond to different preprocessing steps: **(B)** raw beta values, **(C)** beta values preprocessed using SeSaMe’s recommended settings, and **(D)** SeSaMe-preprocessed beta values with adjustment for chamber number batch effects using ComBat.

Notably, while preprocessing with SeSaMe’s recommended pipeline for the EPIC array appeared to center the distributions of beta values, the distribution of beta values for chambers 1 and 8 remained grossly different from the other chambers (Fig 5B-C). Correcting SeSaMe-preprocessed data for chamber number batch effect with ComBat appeared to align the distributions for chambers 1 and 8 such that data from these chambers more closely resembled both data from other chambers as well as a normal distribution (Fig 5D). To evaluate whether similar biases are present between different slides, we similarly converted beta values to within-subject z-scores and averaged scores across samples on the same slide, rather than chamber number (Fig S3A). While biases appear in raw beta values, we noted no gross differences in the distributions after preprocessing with SeSAMe (Fig S3B-C); notably, further adjusting for chamber number biases with ComBat appeared to have no detrimental effect on beta value distributions (Fig S3D).

### Interactions between fluorescence intensity/beta values and chamber number

We next sought to better understand how fluorescence intensities might be impacted by chamber number bias. To ascertain whether chamber number biases were more pronounced in dim or bright probes we examined the difference between measurements from each chamber number from the median across all samples. To draw general conclusions about the relationship between intensity and bias, we binned probes into quantiles by median intensity within each channel. To avoid confounding biological and technical variability, we restricted our analysis to within-subject comparisons.

In raw, SeSAMe-preprocessed, and SeSAMe + ComBat adjusted fluorescence values, we observed that FI appeared to be biased by chamber number, with the magnitude of the bias between chambers appearing to be proportional to the median brightness in both the red and green channels (Fig S4, Fig 6), rather than being limited to only particularly dim or bright probes.

**Figure 6.**
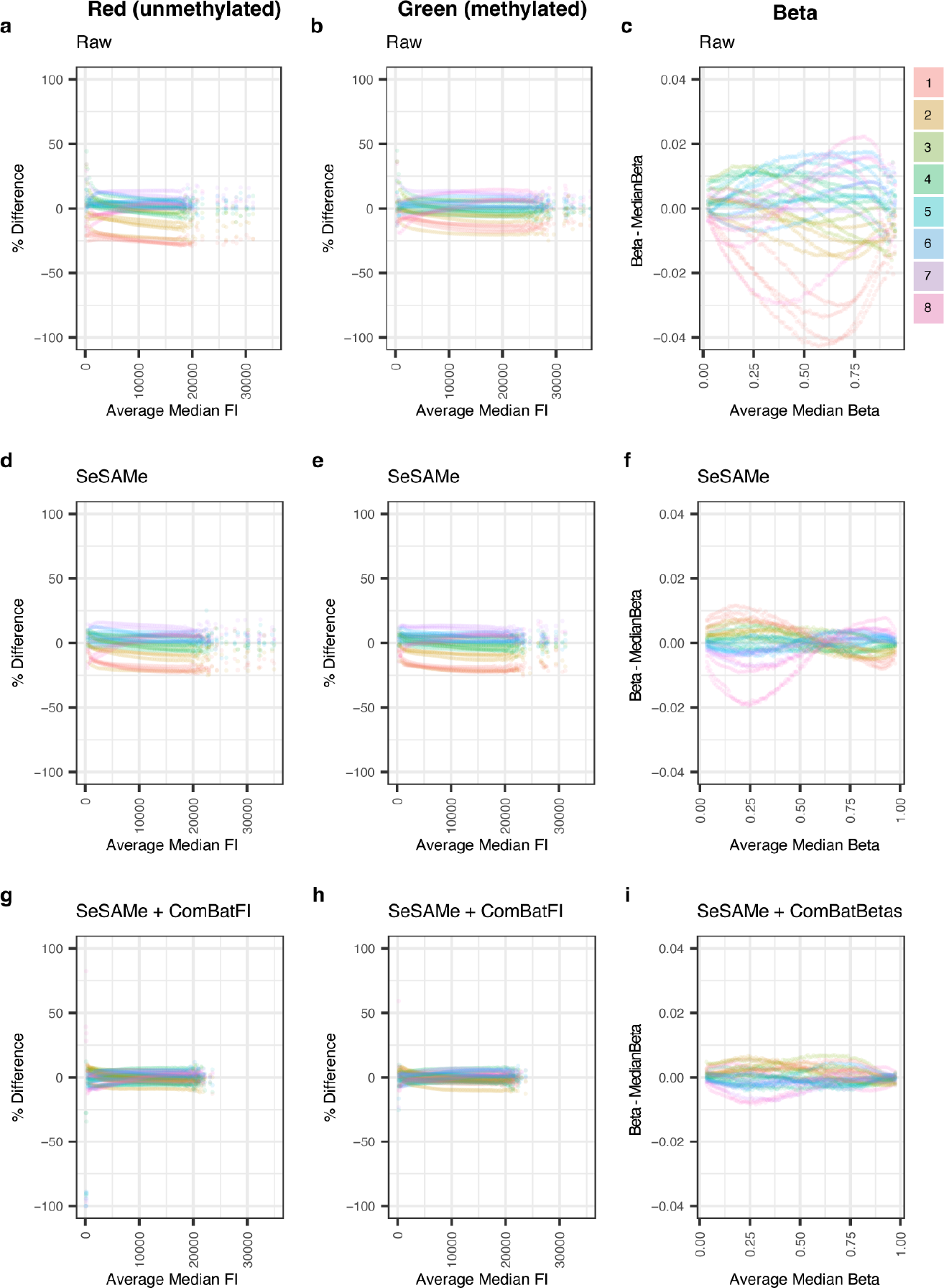
Relative brightness in technical replicates related to probe intensity. Plots depicting Type 2 probes binned according to median within-subject fluorescence intensity (FI) or beta value. The y-axis represents differences from the median value for each bin. Each point shown represents one percentile bin from one of the four subjects in the study. Differences between median intensity are represented for the Red FI, Green FI, and beta values in Raw **(A, B, C)** and SeSAMe-normalized **(D, E, F)** data. Similar comparisons are made after adjusting either the FI **(G-H)** or beta values (I) for Chamber Number using ComBat. Axes for figure panels were intentionally limited, resulting in the omission of some low-intensity data points which had very high variability.

As methylation beta values are directly derived from low-level fluorescence intensities, we next wondered whether beta values are similarly biased in an intensity-dependent fashion. There were pronounced biases in beta values corresponding to the median intensity for intermediate fluorescence intensities; however, these biases appeared to be less consistent than biases within individual color channels (Fig 6C, 6F, 6I).

Importantly, while we observed biases between chamber numbers within each subject, chamber number biases in both fluorescence intensities and beta values appear to be largely consistent across different subjects (Fig S4, Fig 6). Together, these results suggest that positional biases may impact variability and magnitude of probe beta values via chamber-specific differences in probe fluorescence intensity.

### Relationship between variability in low-level data and epigenetic clock predictions

As we had observed intensity-dependent biases with regard to both fluorescence intensity and beta values, we hypothesized that these biases in fluorescence intensity and other low-level data might underlie the variability in beta values. To explore this, we calculated the standard deviation of SeSAMe-preprocessed beta values across all replicate samples for each subject in the study, and then measured the correlation between each of several low-level data features and the standard deviation of beta values (Fig S5). We observed that the number of beads observed for each CpG was inversely correlated with the variability in observed beta values. That is, as the number of beads measuring a CpG increased, the standard deviation of the beta values decreased. Additionally, we observed that the intensity in the green and red channels were also related to variability in beta values, suggesting that data quality might be improved by adjustments for intensity bias.

These results suggest that batch effects and other biases play a substantial role on the overall variability of methylation data, even after common preprocessing methods. However, it remained unclear how or whether such variability is meaningful for common predictive modeling applications using DNA methylation data. To examine this further, we asked whether the variability in predicted ages generated by epigenetic clocks might be explained by variability in low-level data.

To accomplish this aim, we first aggregated predictions from several epigenetic clocks reported in the literature, including Horvath’s clock^15^, Hannum’s clock^16^, Horvath’s Skin and Blood clock^17^, Levine’s PhenoAge clock, and two clocks from Zhang, one trained using best linear unbiased prediction (BLUP) and the other trained using elastic net (EN)^18^. As anticipated, clocks were variable in both the accuracy and precision of predictions (Fig 7D-F). When examining the predicted ages from the Horvath Skin and Blood clock, we noted that age predictions from one technical replicate from subject A stood out as particularly erroneous compared to other technical replicates (Figure 7D). To ascertain what might alter the predicted age from this technical replicate, we examined the variability and coefficients of CpGs within the Horvath Skin and Blood clock. Two CpGs in particular showed a high variability within subject A, in both Raw and SeSAMe-adjusted beta values (Figure 7A-B). We chose to examine one of these, cg03183882, in greater detail due to a larger coefficient within the Horvath Skin and Blood clock (Fig 7A-B). Interestingly, outlier testing indicates that the abnormal technical replicate was a statistical outlier with regards to fluorescence intensity in both the green (Grubbs test for one outlier; p-value < 2.2e-16) and red (Grubbs test for one outlier; p-value = 6.448e-05) channels as well as the standard deviation of fluorescence intensity in green (Grubbs test for one outlier; p-value < 2.2e-16) and red (Grubbs test for one outlier; p-value = 3.269e-10) channels (Fig 7C). Interestingly, adjusting the data with ComBat appears partially compensate for the impact of this CpG on the age predictions of the Horvath Skin and Blood clock (Fig 7D-F). This analysis suggests that CpG outliers detectable within the low-level microarray data may drive variability seen in epigenetic clocks, and that more stringent quality control measures which consider fluorescence measurements might improve microarray performance, as well as the performance of downstream analyses.

**Figure 7:**
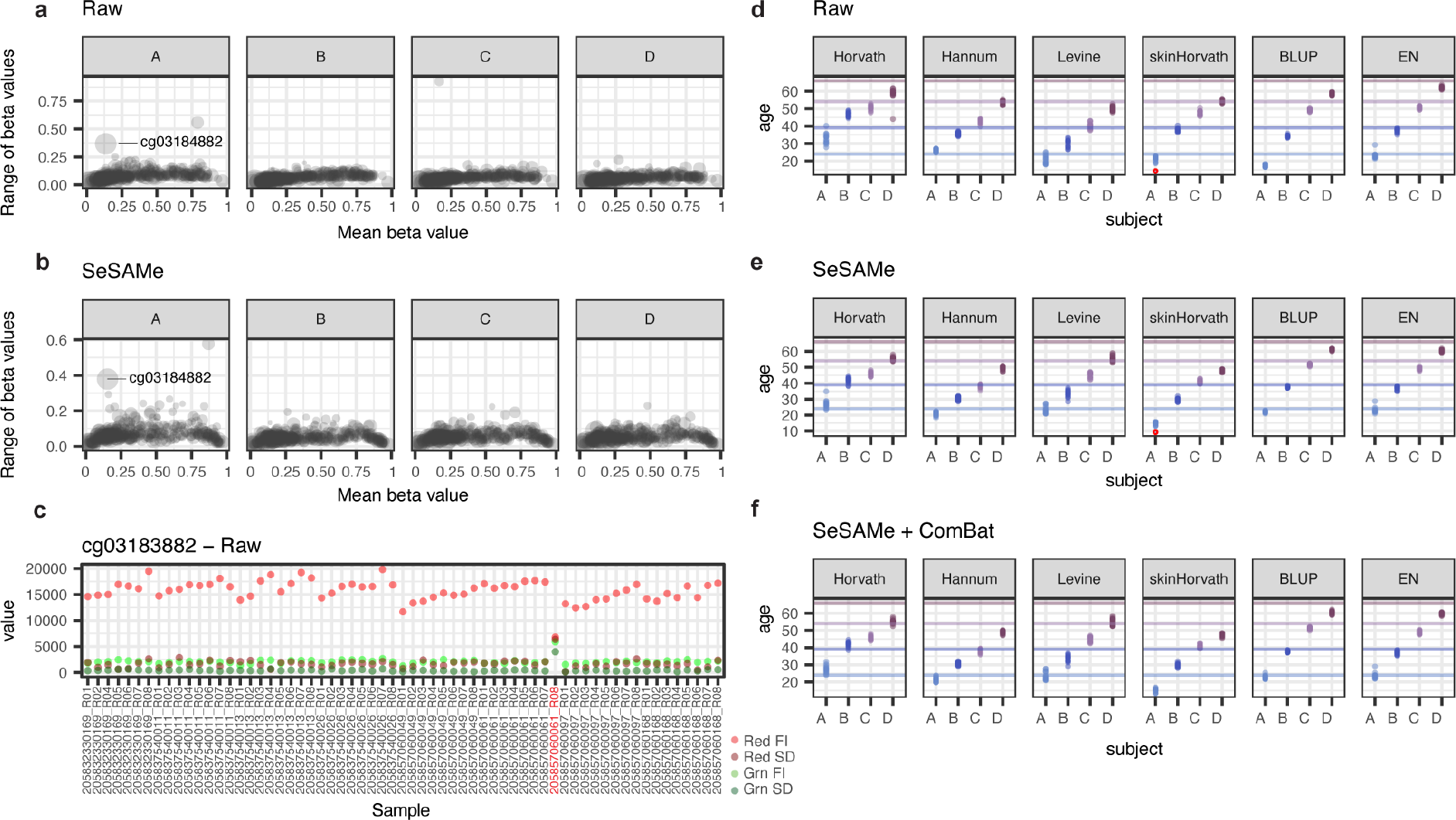
Relationship between variability in low-level data and beta value-derived age-predictions. **(A-B)** Scatter plots displaying the mean vs range of beta values for CpGs included in the Horvath Skin and Blood clock. Each point represents summary statistics from a single CpG across technical replicates from one of the four subjects in the study. Points are scaled according to the coefficient value within the clock model. **(C)** Summary statistics for low-level measurements of cg03184882 within each sample in the study. Outlier sample label is highlighted in red font. **(D-F)** Scatter plots depicting age predictions from 6 different DNAm clocks on technical replicates. Colors of dots and lines indicate predicted age and actual age of subjects, respectively. Red point indicates outlier sample highlighted in panel C. **(D)** Age predictions generated using raw beta values. **(E)** Age predictions generated using beta values generated using the ‘recommended’ preprocessing settings from SeSaMe. **(F)** Age predictions generated using SeSaMe-preprocessed beta values with chamber number batch effect correction using ComBat.

## Discussion

In this study, we explore variability in the Illumina MethylationEPIC microarray data using several highly similar technical replicates, allowing us to isolate and measure the technical variability arising from positional effects within the MethylationEPIC v1 platform in a way that is, to our knowledge, unique within the published literature. We assume that all variance observed between pooled samples arises from technical noise or biases within the microarrays themselves, rather than any biological variability. Similarly, due to the aggregate nature of measurements using Illumina BeadChip technology, with fluorescence intensity measurements being derived from the fluorescence of many probes per bead, and these also averaged across usually several beads per CpG, we discount the idea that sampling bias might play a meaningful role in observed differences between technical replicates.

First, using principal compenents analysis, we demonstrate the occlusion of a strong biological signal by positional effects within the array. In this study, we generated technical replicates with a far higher similarity than might be reasonably expected from any experiment. Despite this, the chamber number positional effects on the samples were sufficient to result in poor hierarchical clustering of replicates from the same subject within PCA space until corrected for using ComBat. This finding has notable implications for previous studies of microarray reliability using technical duplicates.

Next, using linear mixed effect models, we quantified variability, distinguishing the overall contributions of slides and chamber numbers to the variance of methylation beta values. These inescapable positional effects are present even within small microarray experiments, and numerous methods have been developed to correct these effects. While many publications have explored these batch effects and make recommendations regarding the balancing of experimental groups across arrays and chamber numbers, the present study is the first, to our knowledge, to quantify the magnitude of these batch variables.

Our observations suggest that in raw data, the chamber number positional effect has a larger overall effect than slide on beta value variability; however, the effect of chamber number appears to be better corrected than slide effects using existing preprocessing tools such as the ComBat algorithm. This finding suggests that different balancing strategies should be utilized, depending on the analysis pipeline being used, and has immediate implications for the researcher attempting to minimize the prevalence of experimental errors due to positional effects: if the experiment is sufficiently large to allow for chamber number correction using ComBat, then prioritization should be given to balancing experimental groups across slides; and if not, then balancing groups across chamber numbers should be given priority.

Previous efforts at improving the reliability of DNA methylation microarrays have used varying approaches, including building clocks based off of principal components rather than individual CpGs. Our analyses suggest that these approaches, in the absence of appropriate preprocessing, may confuse technical biases in the data with biological signal, and may partially explain why many epigenetic clocks tranfer poorly to other datasets. Even within highly similar technical replicates and widely used preprocessing tools, we observed that positional effects from the microarray were highly represented in principal components and obfuscated biological variability. However, we note that after correcting beta values with ComBat, principal components become largely driven by biological differences between samples.

By exploring low-level data, we observed that positional effects appear to be largely proportional to total fluorescence intensity, with batch effects most apparent in those probes with either very dim or very bright fluorescence intensities (combined red and green channels). While notable, these effects appear to be difficult to predict, and are inconsistent both between subjects and between slides. Further, we observed the presence of technical outliers within low-level data which appear to escape any preprocessing and may impact the performance of epigenetic clocks. We anticipate that increasing the stringency of outlier filtering would likely result in increases in performance; however, the most appropriate method of detecting and filtering such outliers remains unexplored.

We note in this manuscript the apparent benefits of preprocessing with SeSAMe as well as chamber number batch adjustment with ComBat for several metrics of variability and that these processing step dramatically reduces the number of false positive findings on differential methylation tests, while preserving detectable biological variability and improving the consistency of measurements.

We also explored an observation of a technical outlier for a single CpG with a high coefficient in methylation age clock predicitons. This observation appears to be an outlier with regards to both fluorescence intensity and variability in both red and green channels, and these data were recorded from relatively few beads (nBeads = 4). ComBat appeared to improve the precision of biological age predictions, including adjusting this outlier to be more similar to other technical replicates; however, such outlier values among technical replicates suggests an unmet need for quality control methods that consider within-sample variability in low-level fluorescence measurements, perhaps utilizing the existing standard deviation measurements within BeadChip technology.

Finally, we note that ComBat does not reduce variability in the epigenetic-clock predicted ages of our technical replicates. However, existing epigenetic clocks such as those tested here were trained with little regard for positional effects. While such effects might be partially controlled for by the machine learning methods themselves, the optimization of preprocessing methods for the development of computational models remains largely unexplored.

## Acknowledgements

We would like to acknowledge Diagenode and BioIVT for assisting with sample processing and curation of the dataset. We would also like to acknowledge Wanding Zhou for helpful review prior to submission.

## Author contributions

AB conceived of experimental methods, performed experiments, collected results, and wrote the manuscript. JK conceived experimental methods and collected preliminary results. ER performed initial preprocessing of microarray data. IB conceived experimental methods. DLV designed initial experiments, conceived experimental methods, supervised the project, and assisted in revising the manuscript. KC generated datasets, assisted with experimental design, and helped guide the overall study. DV conceived experimental methods and supervised the project. All authors discussed results and contributed to the final manuscript.

## Materials and Methods

### Sample Collection

Whole blood samples from four adult males were collected by and purchased from BioIVT. At the time of enrollment, all individual donors provided informed consent that their samples and genetic information could be used for future studies, including this study. All identifying clinical information was removed from samples except for approximate age and ethnicity. All donors were screened and found to be hepatitis B surface antigen negative, HBV NAT negative, HIV 1&2 antibody negative, HIV NAT negative, HCV antibody negative, HCV NAT negative, syphilis negative, West Nile virus NAT negative, and certified to have been negative for antibodies to *T. cruzi* either on the current or at least one previous donation. CBCs were performed and 150?l of blood was aliquoted into 1.5mL tubes and frozen on dry ice for direct shipping to Diagenode.

### DNA Extraction and Microarray Assay

Total DNA was extracted from eight 150 μl aliquots of anticoagulated blood per subject using DNeasy Blood and Tissue Kit (Qiagen, Cat No. 69504) with RNase treatment option, using RNAse cocktail from Thermofisher (AM2286). Final elution of DNA was carried out in 200μl of elution buffer, and samples were quantified using Qubit dsDNA HS Assay Kit (Thermo Fisher Scientific). DNA quality was assessed using the Fragment Analyzer and DNF-488 High Sensitivity genomic DNA Analysis Kit (Agilent). Concentrations and Fragment Analyzer profiles are included in supplemental materials.

Each of the resulting eight DNA aliquots per subject was further split into two aliquots, 64 in total, and independently deaminated, amplified, and fragmented. Deamination was carried out using EZ-96 DNA Methylation Kit (Zymo Research) according to Illumina’s recommended deamination protocol. An aliquot of DNA from each of eight technical replicates was combined into a pooled DNA sample for each subject. A total of eight independently isolated replicates for each subject and eight aliquots of the pooled sample for each subject were measured using Illumina Infinium MethylationEPICv1 array BeadChips (850K). The above steps were carried out by the Epigenomic Services from Diagenode (Cat nr. G02090000).

### Data availability

Data generated in this study are available in the Gene Expression Omnibus (GEO) and accessible through GEO series number GSE250556. These data include raw microarray data files and normalized methylation data generated using Illumina MethylationEPICv1 arrays. Code used in data processing, analysis, and generation of results are available at the following GitHub repository: https://github.com/AndersonButler/EPICv1_variability.

### Data preprocessing

Raw data in the form of idat files were loaded and preprocessed using the SeSAMe, R package to obtain either raw fluorescence or beta values (prep = ““) or fluorescence/beta values calculated using the recommended settings (prep = “QCDPB”) using the built-in manifest^19^.

To obtain ComBat-adjusted beta values, SeSAMe-adjusted values were then further preprocessed by removing CpGs with greater than 10% NA values among all samples. Missing values among remaining CpGs were median imputed. This additional step resulted in a total 614,831 CpGs which were shared across all preprocessing pipelines discussed above. Values were then adjusted using ComBat from the sva R package. To obtain ComBatFI-adjusted beta values, SeSAMe-adjusted fluorescence intensities were adjusted using the ComBat-Seq function from the sva package.

A single technical replicate from the pooled samples was removed as an outlier based on quality (205832330169_R03C01, subject C). After SeSAMe preprocessing, one technical replicate had an abnormally low rate of successfully detected probes (Grubbs test for outlier, p<0.05). This sample was removed from all subsequent analyses.

Where indicated, fluorescence values or beta values for each CpG were converted to z-scores by centering and scaling, by subtracting the within-subject mean and dividing by the within-subject standard deviation among technical replicates using the scale function in R. Z-score analyses were filtered to include only CpGs with values complete and passing quality thresholds in >90% of samples. CpGs with non-finite z-score converted values were omitted from plots.

### Statistical analyses

#### Linear mixed effects models

Beta values from raw data, SeSAMe-preprocessed data, or SeSAMe-preprocessed data adjusted with ComBat for chamber number batch effects were cleaned by removing CpGs with greater than 10% NA values among all samples and median imputing the remaining missing values as described above. Linear mixed effects regression was conducted using the lmer function from the lme4 R package, using beta values as the dependent variable, slide and chamber numbers as fixed effects, and subject as a random effect. Analysis of variance (ANOVA) was conducted on models using the F-test, to assess the importance of positional effects in explaining the variation in the beta values.

#### Principal components analysis and hierarchical clustering

Beta values from SeSAMe-preprocessed data, or SeSAMe-preprocessed data adjusted with ComBat for either chamber number or slide number as batch effects were further preprocessed by removing CpGs with greater than 10% NA values among all samples, missing values among remaining CpGs were median imputed from all samples. The first 50 principal components were calculated using the factoextra R package (v1.0.7). For hierarchical clustering, the hclust function was used with Ward’s D2 method.

#### Differential methylation analysis

Differential methylation testing for each set of comparison groups was conducted using the dmpFinder function from the minfi R package. Note that for two-comparison tests, the results are similar to Student’s t-test. Multiple comparisons corrections were conducted using the Benjamini-Hochberg method.

#### Standard deviation ratios

Ratios of the standard deviations for within-subject and across subject comparisons were calculated by averaging the standard deviations of technical replicates for each subject and dividing this average by the standard deviation of all replicates across all subjects with the addition of a small offset of 0.0001 to account for low variability CpGs.

#### Correlation plots

Low level data were extracted from idat files using the R packages UMtools^20^ and Watermelon^21^. Correlation matrices were calculated and plotted using the R package corrplot.

### Epigenetic clocks

Age predictions from epigenetic clocks were calculated using the methylclock R package for the Horvath, Hannum, Horvath Skin and Blood, Levine, BLUP, and EN clocks^22^.

**Figure S1.**
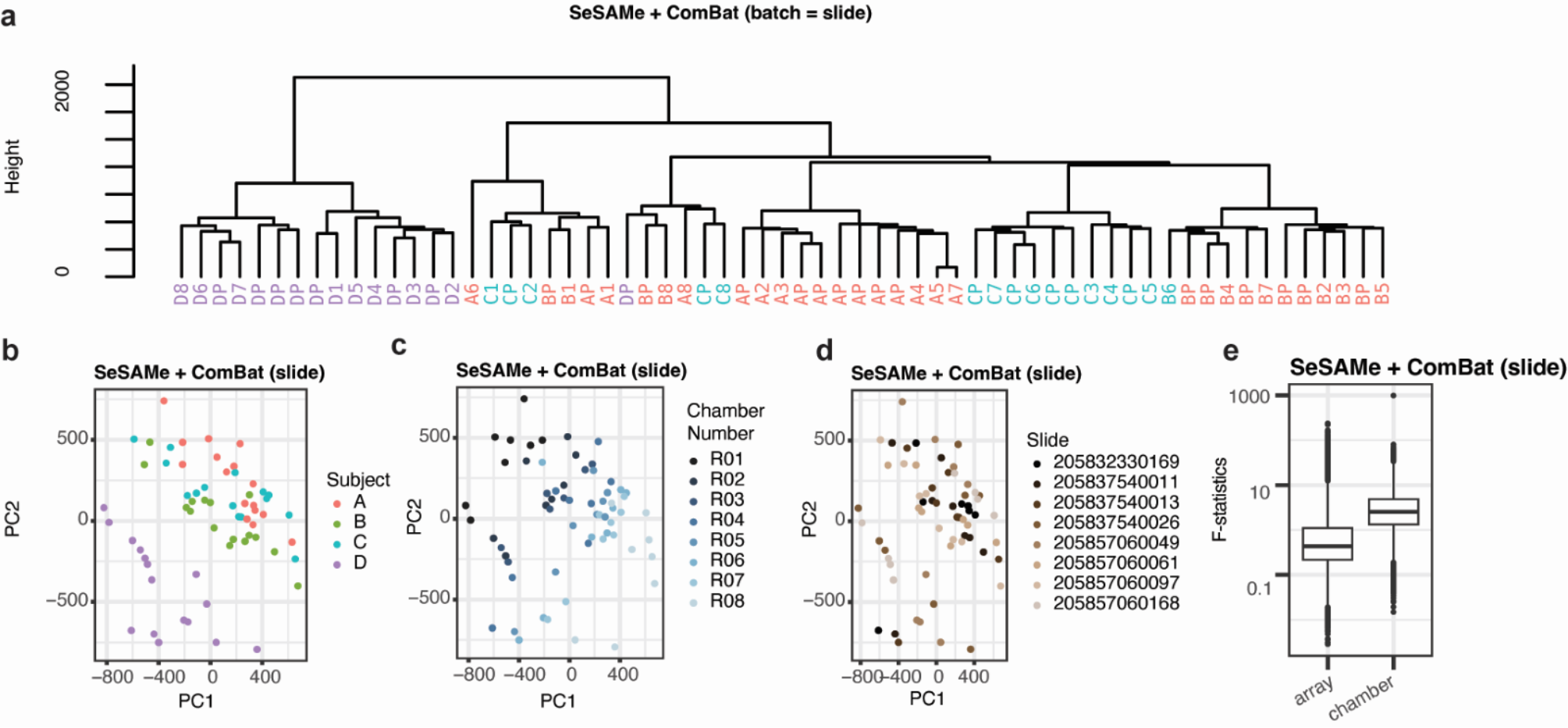
PCA and Hierarchical Clustering of Beta Values After ComBat using Slide as Batch Effect. **(A)** Hierarchical clustering dendrogram representing the clustering of subjects based on the first 50 principal components from adjusted beta values after preprocessing with SeSAMe and ComBat adjustments for slide. **(B-D)** PCA plots showing clustering of subjects after ComBat adjustment by slide, colored by subject **(B)**, chamber number **(C)**, or array **(D). (E)** Box plot presenting F statistics obtained from analysis of variance (ANOVA) performed on beta values preprocessed with SeSAMe + ComBat using slide as batch. The box represents the interquartile range (IQR), with the median indicated by a line inside the box. Whiskers extend to the minimum and maximum values within 1.5 times the IQR. Data points beyond 1.5 times the IQR are plotted individually.

**Figure S2.**
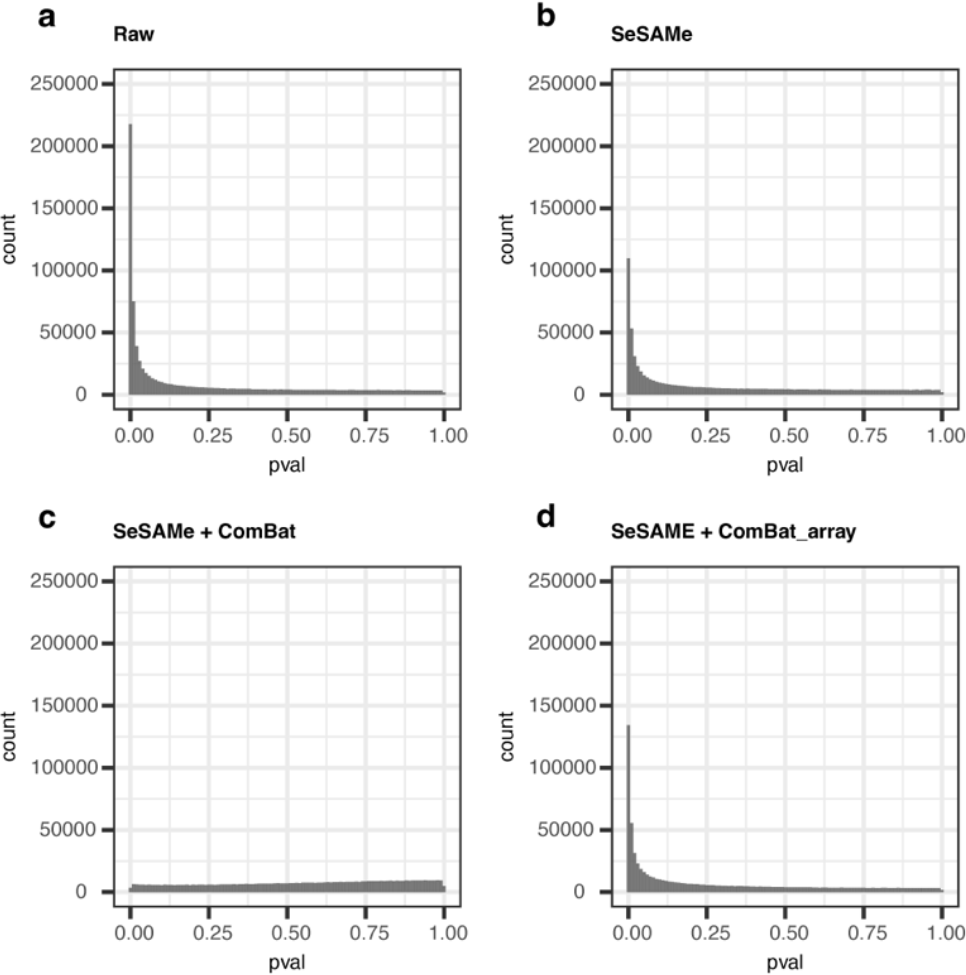
Differential methylation analysis of alternatively preprocessed methylation beta values from technical replicates without FDR correction. **(A-D)** Histograms illustrate distribution of uncorrected p-values obtained from probewise differential methylation testing on raw beta values **(A)**, beta values preprocessed using SeSaMe’s recommended settings **(B)**, and **(C-D)** SeSaMe-preprocessed beta values with correction for batch effects associated with chamber number using ComBat to adjust for either chamber number **(C)** or slide **(D)**. times the IQR are plotted individually.

**Figure S3:**
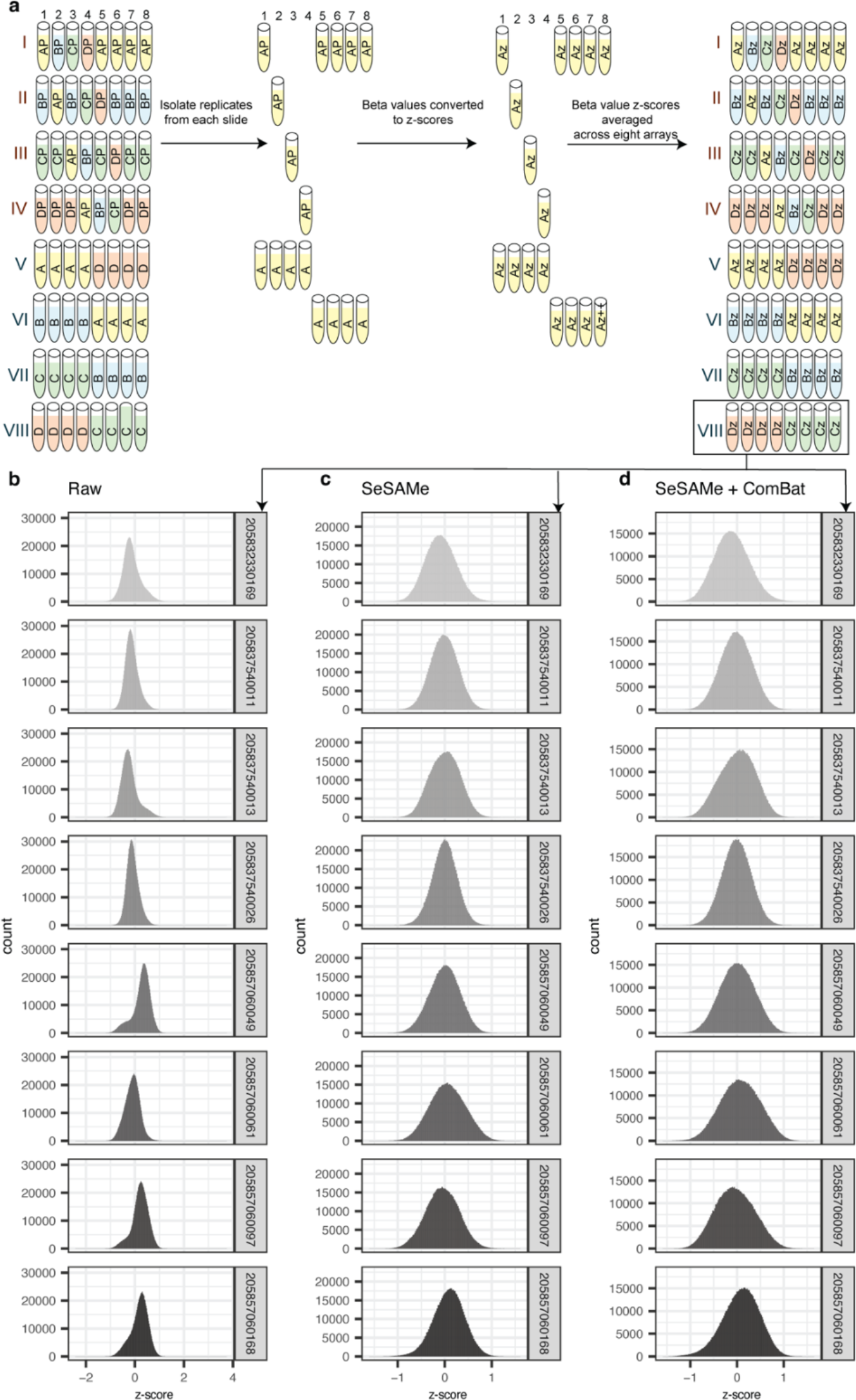
Beta values compared in technical replicates measured in different slides. **(A)** Configuration of samples within slides/chambers. The average of within-subject centered and scaled beta values for each CpG/slide were extracted from all chamber numbers on each slide as depicted. **(B-D)** Histograms representing the distributions of within-subject centered and scaled beta values for all probes on the MethylationEPIC array. The histograms correspond to different preprocessing steps: **(B)** raw beta values, **(C)** beta values preprocessed using SeSaMe’s recommended settings, and **(D)** SeSaMe-preprocessed beta values with adjustment for chamber number batch effects using ComBat. All panels were derived from eight samples except array 205832330169, from which one sample was discarded due to low quality.

**Figure S4.**
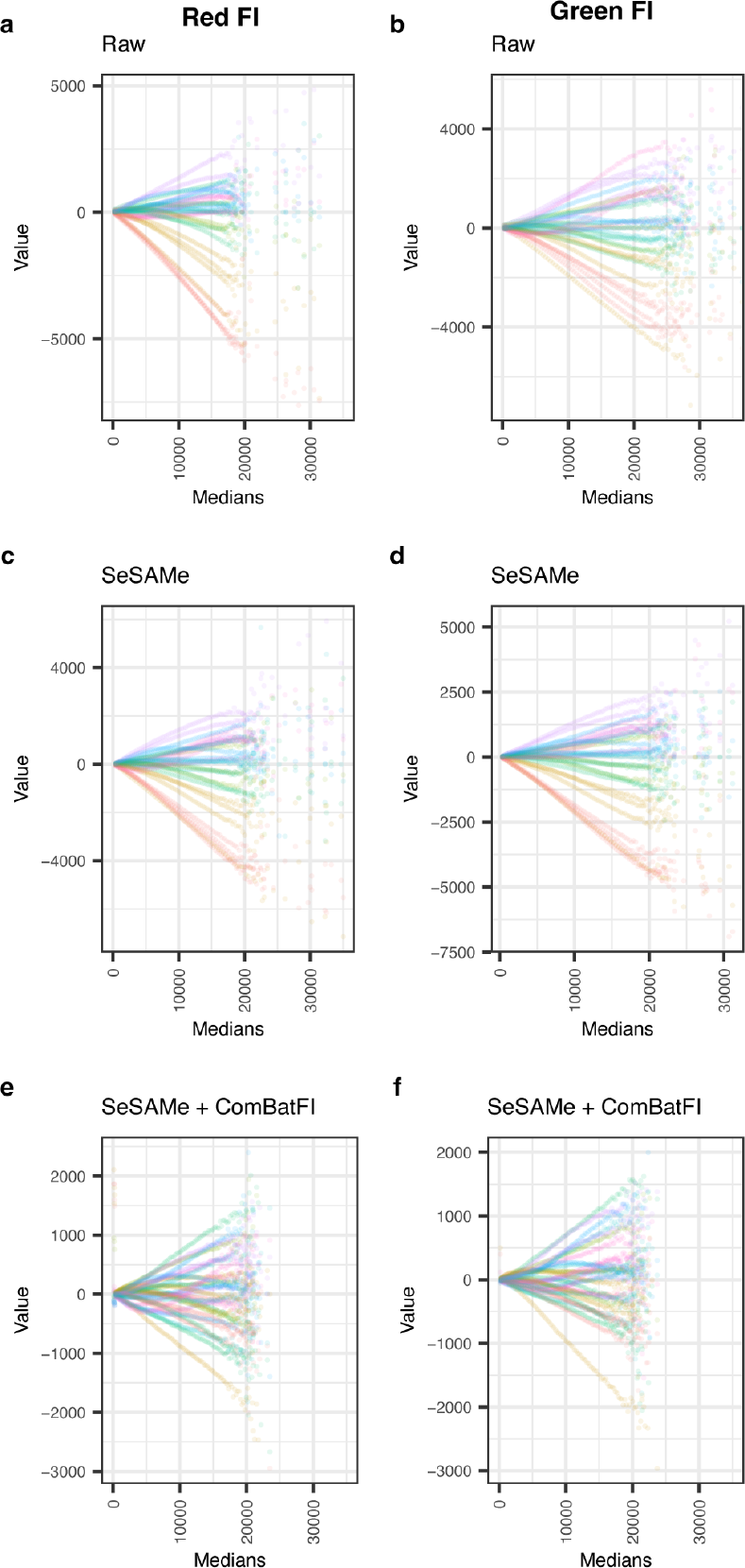
Relative brightness in technical replicates is correlated with probe intensity. **(A-F)** Plots depicting the difference from the median in percentile bins, scaled in fluorescence intensity units. Each point shown represents one percentile bin from one of four subjects. Raw, SeSAMe-normalized, and ComBat-corrected values are shown for the green **(a, c, e)** and red **(b, d, f)** channels, respectively.

**Figure S5:**
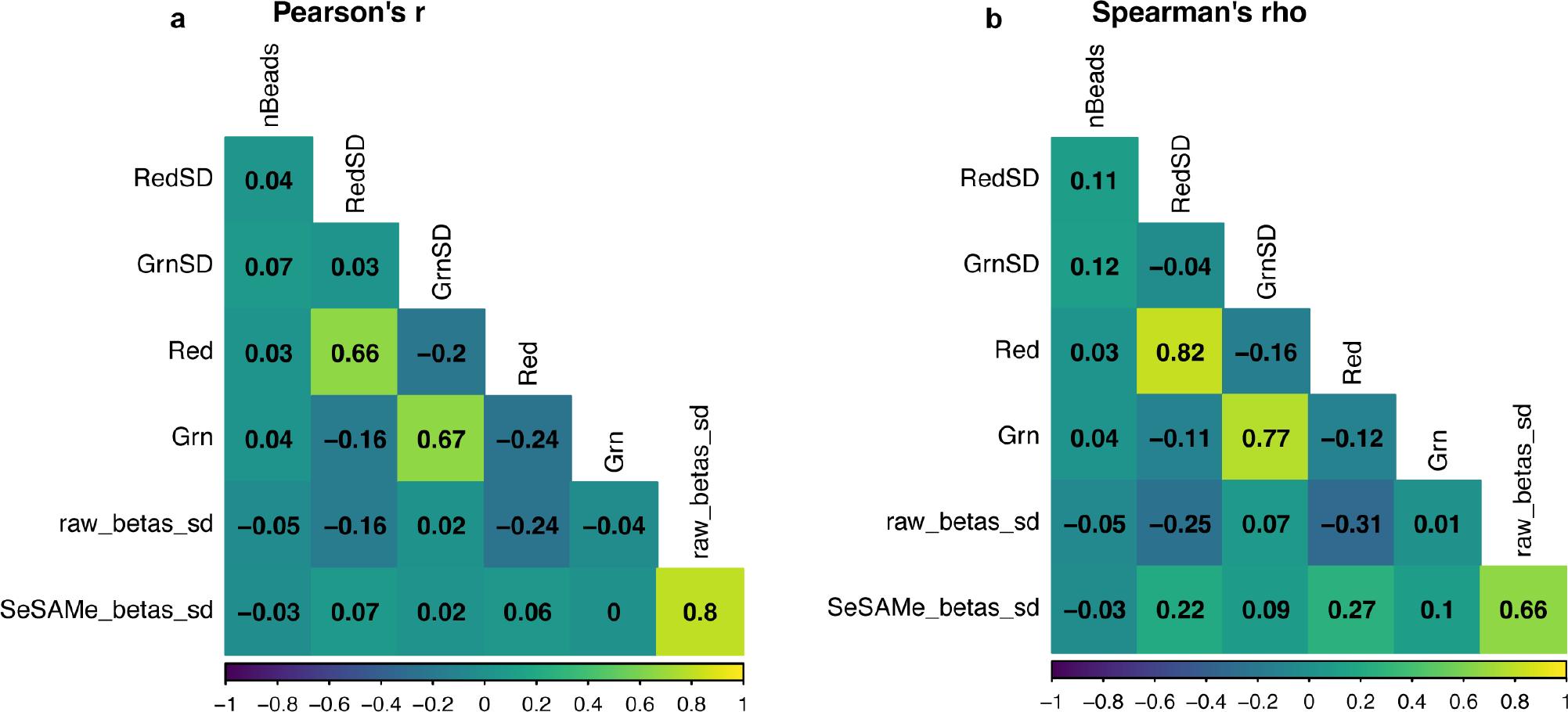
Relationship between low-level variables and beta values. Plot depicts Pearson’s R (A) and Spearman’s rho (B) calculated for low-level variables and the standard deviation (SD) of either raw or SeSAMe-preprocessed beta values of Type 2 probes. Only probes present in both raw and Sesame-preprocessed data were used to calculate correlations.

## Bibliography

1. Lu, A. T. et al. DNA methylation GrimAge strongly predicts lifespan and healthspan. Aging Albany Ny 11, 303–327 (2019).

2. Lu, A. T. et al. DNA methylation GrimAge version 2. Aging (Albany NY) 14, 9484–9549 (2022).

3. Nabais, M. F. et al. An overview of DNA methylation-derived trait score methods and applications. Genome Biol. 24, 28 (2023).

4. Moran, S. et al. Epigenetic profiling to classify cancer of unknown primary: a multicentre, retrospective analysis. Lancet Oncol. 17, 1386–1395 (2016).

5. Taryma-Lesniak, O., Sokolowska, K. E. & Wojdacz, T. K. Current status of development of methylation biomarkers for in vitro diagnostic IVD applications. Clin. Epigenetics 12, 100 (2020).

6. Walter, S. D., Eliasziw, M. & Donner, A. Sample size and optimal designs for reliability studies. Stat. Med. 17, 101–110 (1998).

7. Watson, P. F. & Petrie, A. Method agreement analysis: A review of correct methodology. Theriogenology 73, 1167–1179 (2010).

8. Xu, Z. & Taylor, J. A. Reliability of DNA methylation measures using Illumina methylation BeadChip. Epigenetics 16, 495–502 (2021).

9. Dugué, P.-A. et al. Reliability of DNA methylation measures from dried blood spots and mononuclear cells using the HumanMethylation450k BeadArray. Sci Rep-uk 6, 30317 (2016).

10. Sugden, K. et al. Patterns of Reliability: Assessing the Reproducibility and Integrity of DNA Methylation Measurement. Patterns 1, 100014 (2020).

11. Logue, M. W. et al. The correlation of methylation levels measured using Illumina 450K and EPIC BeadChips in blood samples. Epigenomics-uk 9, 1363–1371 (2017).

12. Higgins-Chen, A. T. et al. A computational solution for bolstering reliability of epigenetic clocks: implications for clinical trials and longitudinal tracking. Nat Aging 1–18 (2022) doi:10.1038/s43587-022-00248-2.

13. Zindler, T., Frieling, H., Neyazi, A., Bleich, S. & Friedel, E. Simulating ComBat: how batch correction can lead to the systematic introduction of false positive results in DNA methylation microarray studies. Bmc Bioinformatics 21, 271 (2020).

14. Zhang, Y., Parmigiani, G. & Johnson, W. E. ComBat-seq: batch effect adjustment for RNA-seq count data. Nar Genom Bioinform 2, qaa078-(2020).

15. Horvath, S. DNA methylation age of human tissues and cell types. Genome Biol 14, R115–R115 (2013).

16. Hannum, G. et al. Genome-wide Methylation Profiles Reveal Quantitative Views of Human Aging Rates. Mol. Cell 49, 359–367 (2013).

17. Horvath, S. et al. Epigenetic clock for skin and blood cells applied to Hutchinson Gilford Progeria Syndrome and ex vivo studies. Aging Albany Ny 10, 1758–1775 (2018).

18. Zhang, Q. et al. Improved precision of epigenetic clock estimates across tissues and its implication for biological ageing. Genome Med 11, 54 (2019).

19. Zhou, W., Triche, T. J., Laird, P. W. & Shen, H. SeSAMe: reducing artifactual detection of DNA methylation by Infinium BeadChips in genomic deletions. Nucleic Acids Res 46, gky691. (2018).

20. Jiménez, B. P., Kayser, M. & Vidaki, A. Revisiting genetic artifacts on DNA methylation microarrays exposes novel biological implications. Genome Biol 22, 274 (2021).

21. Pidsley, R. et al. A data-driven approach to preprocessing Illumina 450K methylation array data. Bmc Genomics 14, 293 (2013).

22. Pelegí-Sisó, D., Prado, P. de, Ronkainen, J., Bustamante, M. & González, J. R. methylclock : a Bioconductor package to estimate DNA methylation age. Bioinformatics 37, 1759–1760 (2020).

